# FRC-QE: A robust and comparable 3D microscopy image quality metric for cleared organoids

**DOI:** 10.1101/2020.09.10.291286

**Authors:** Friedrich Preusser, Natália dos Santos, Jörg Contzen, Harald Stachelscheid, Érico Tosoni Costa, Philipp Mergenthaler, Stephan Preibisch

## Abstract

Three-dimensional stem-cell-derived organoids are a powerful tool for studying cellular processes in tissue-like structures, enabling in vitro experiments in an organ-specific context. While organoid research has been closely linked to advances in fluorescence microscopy, capturing cellular structures within their global context in an organoid often remains challenging due to the organoid’s dense structure and opacity. The development of optical clearing methods has provided a solution for fixed organoids but optimizing clearing protocols for a given sample type and staining can be challenging. Importantly, quantitative measures for assessing image quality throughout cleared fluorescent samples are missing. Here, we propose Fourier ring correlation quality estimation (FRC-QE) as a new metric for automated 3D image quality estimation in cleared organoids. We show that FRC-QE robustly captures differences in clearing efficiency within an organoid, across replicates and clearing protocols, as well as for different microscopy modalities. FRC-QE is open-source, written in ImgLib2 and provided as an easy-to-use and macro-scriptable plugin for the popular Fiji software. We therefore envision FRC-QE to fill the gap of providing a reliable quality metric for testing, optimizing and comparing optical clearing methods.

## Introduction

Organoid models have emerged as powerful tools to investigate fundamental biological questions. Indeed, human organoid models in general offer the opportunity to investigate complex biological processes or personalized therapies in a human model system in an organ-specific context (1, 2). While organoid models have increased in complexity and single cell analysis techniques are commonly applied to them, quantitative three-dimensional (3D) microscopy of these models at single-cell resolution is a difficult task. In particular, size and opacity of these complex tissuelike structures represent a major limitation for acquiring 3D microscopy images. Since its inception (3), tissue clearing has emerged as a powerful method that enables imaging of opaque, large, fixed samples with single-cell resolution (4–6). However, establishing an optimal clearing pipeline for a particular sample, staining, and microscopy setup remains challenging.

Experimentalists can choose from a large variety of different clearing protocols that differ in terms of experimental complexity, reproducibility, time, cost, flexibility, compatibility with staining methods and imaging setups, as well as, most importantly, clearing performance on the sample of interest (7). Choosing the most suitable protocol therefore requires unbiased comparison of clearing performance. However, while significant efforts are put into the development of new clearing methods, the current state-of-the-art for assessment of clearing performance is typically manual inspection of light diffraction through the sample using a printed raster image. While this represents a simple way of assessing global sample transparency, it does not yield a quantitative measurement and is not specific for a fluorescent signal of interest. Moreover, the optical set-up used for this type of images is different from a fluorescence microscope (e.g. light-sheet or spinning-disk confocal) that will be used to perform 3D fluorescence imaging. Hence, assessing clearing efficiency and final image quality based on the visibility of the raster image does not reflect the actual experimental set-up and will not take into account potential pitfalls arising in fluorescence microscopy (e.g. increased autofluorescence or bleaching of the cleared sample). Furthermore, it does not contain spatial information regarding the cleared sample, i.e. how far into the sample can fluorescent structures of interest be resolved in the case of insufficient clearing. Alternatively, manual inspection of fluorescence images is very time consuming, not quantitative and thus makes comparison in between different protocols difficult. To obtain a quantitative measure of image quality, statistics such as intensity or contrast (i.e. relative intensity) of the sample are sometimes used to provide a quantitative read-out (8–10) of clearing efficiency. However, they do not necessarily reflect actual image quality, and are not comparable across protocols or microscopes, often not even between experimental replicates. To address these issues, we propose the Fourier ring correlation quality metric (FRC-QE) that provides a robust readout of image quality for three-dimensional samples (Figure 1). We demonstrate the power of FRC-QE by solving the challenge of identifying a suitable clearing protocol and microscopy setup for brain organoid imaging.

**Fig. 1.**
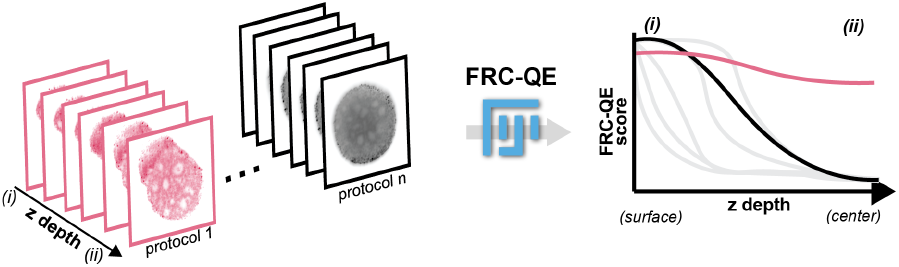
Fourier ring correlation quality estimate for clearing efficiency. Clearing efficiency of multiple samples can differ across experiments due to different protocols or experimental variability. Here, the sample cleared with protocol n has decreased image resolution in the center region (ii) compared to the surface region (i). Our Fourier ring correlation quality estimate (FRC-QE) plugin is designed to automatically quantify clearing efficiency across multiple protocols and is provided as a FIJI plug-in.

## Results

### FRC-QE reflects image quality across cleared organoids

To test whether FRC-QE successfully recapitulates image quality throughout an organoid sample, we first compared FRC-QE to established image quality metrics: Intensity, contrast and also Shannon entropy that has not been proposed in this context yet (Fig. 2). For comparison between organoid replicates, we generated human induced pluripotent stem cell (hiPSC)-derived cerebral organoids of a defined size (ca. 600 μm) that were stained and subsequently chemically fixed. We used the fluorescent DNA stain Draq5 as reference for image quality estimation because (i) it is a small molecule diffusing through organoids independent of clearing efficiency (11), (ii) it stains nuclei, a cellular compartment that is homogeneously located throughout the organoid and (iii) because its fluorescence spectrum is in the farred spectrum (excitation: 647 nm, emission: 696 nm), resulting in better light penetrance through dense tissue when compared to dyes with lower wavelengths (e.g. DAPI). To compare image quality across whole organoids between different clearing methods, we chose three published and straight-forward to implement clearing methods as proof of concept and applied them to our samples (Supplementary figure 1). We used ClearT2 (12), ScaleA2 (13), and fructose-glycerol (14) clearing, which vary with regard to their experimental complexity and time. Cleared organoids were then imaged by multi-view light-sheet microscopy (15) and reconstructed computationally (16). Since organoids were imaged from multiple angles and illumination directions, light scattering is increased in the middle of the organoid compared to its surface. We therefore expected increased resolution of cellular features at the edges and decreased image quality towards the center of the organoid. Homogeneous image quality across the entire sample (e.g. surface vs. center of the organoid) would indicate the absence of light scattering and thereby perfect clearing. Figure 2a-d shows sample images of an insufficiently cleared organoid. To quantify this effect, we processed a 400×400 pixel volume within the center of the organoid, spanning all z-slices of the entire volume (Figure 2a-d, enlarged section), which we used for all following analyses (Figure 2e-h). While individual nuclei were clearly resolved at the edges of the organoid (Figure 2b,d), the center region remained blurred (Figure 2c), indicating increased light scattering caused by insufficient clearing. Importantly, the obvious decrease in image quality towards the center was not faithfully captured when quantifying pixel intensity across the z-axis (Figure 2e). The intensity measurement is correlated with the amount of fluorescent structures within the field of view at a given z-position and does not contain reflect the image quality of the resolved features. Alternatively, image contrast measures the difference between resolved features and background, therefore potentially containing information about the signal-to-noise ratio and thus image quality. While measuring contrast across the z-axis does indeed capture the improved resolution at the surface of the organoid, it is sensitive to artifacts (Figure 2f, peaks in edge region).

**Fig. 2.**
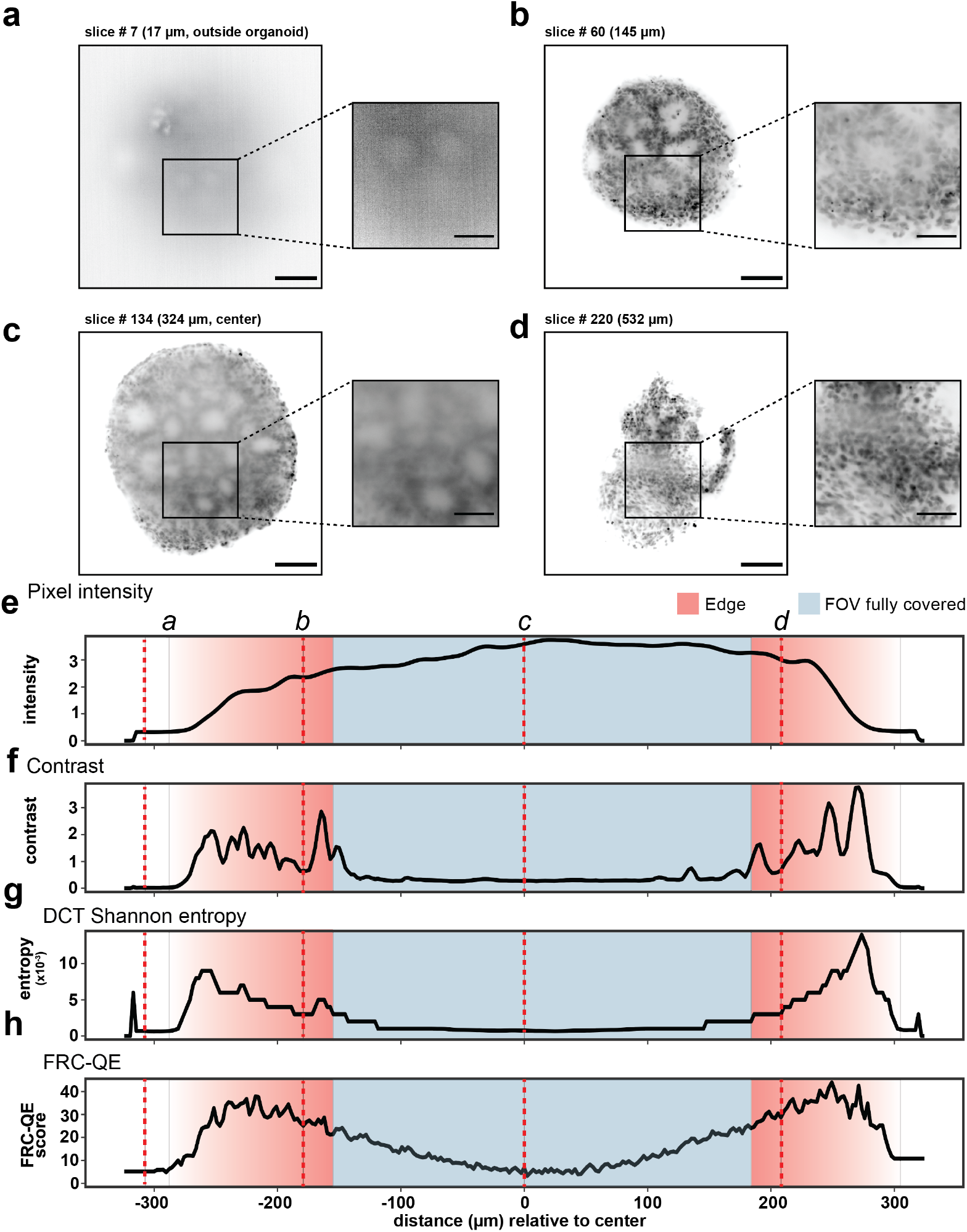
Quantification of the image quality throughout the organoid. (a-d) Optical sections throughout an organoid cleared with the ScaleA2 protocol, stained with Draq5 and reconstructed from a multi-view light-sheet acquisition. Image resolution decreases towards the middle of the organoid as seen in (c). (e-h) Different quantification modalities to assess image quality of the same organoid depicted above. Dotted red lines correspond to the corresponding panels above as indicated. Scale bars correspond to 100 μm and 50 μm for large panels and inlets, respectively. Brightness and contrast was adjusted individually for the example images.

Notably, at the organoid surface, where pixel intensity differs significantly between cells and background, contrast fluctuates and can decrease to values that are closer to the values measured at the center of the organoid. However, image quality is worse within the center than at these surface regions (Figure 2c). Therefore, the contrast curves do not reflect clearing efficiency. Moreover, the dense center of the organoid has a very low signal-to-noise ratio, resulting in a contrast value close to zero throughout the majority of z-slices. Hence, the resulting low dynamic range does not enable quantification of the observed differences in image quality that occur deeper in the organoid. Since both image intensity and contrast measurements did not sufficiently recapitulate image quality, we calculated the normalized discrete cosine transform (DCT) Shannon entropy for each slice across the z-axis. Shannon entropy of the DCT measures the information content of the frequency space and has been previously used for assessing image quality of fluorescence images, notably in the context of automated microscopy (17, 18). DCT Shannon entropy captures the relative difference in image quality across the organoid showing increased entropy at the surface compared to the center region (Figure 2g), thereby outperforming plain intensity and contrast measurements. Fourier ring correlation (FRC) has been previously used to measure resolution in both electron (19, 20) and fluorescence microscopy (21–23). We previously extended this concept to three-dimensional fluorescence microscopy (16) by approximating independent observations using subsequent slices in a stack of the same object and normalization to more distant slices that we further adjusted here (see methods). With this method that we call Fourier ring correlation quality estimate (FRC-QE), we are able to assess image quality across the z-axis of the organoid. To test whether FRC-QE sufficiently describes relative differences in image quality across the z-axis, we analyzed the same pixel volume as before. A relative FRC score was calculated for each slice within that subvolume (Supplementary figure 2). Based on our measurements we conclude that FRC-QE does indeed capture clearing efficiency for three reasons: (i) The FRC-QE score across the organoid reflects the increased resolution at the surface of the organoid (Figure 2h, red shaded areas). (ii) The measurement is less prone to artifacts, e.g. compared to the contrast measurement, resulting in a smooth curve throughout the z-axis. (iii) FRC-QE disposes of the needed dynamical range to quantify differences in clearing efficiency even with increased light scattering (i.e. in the center of the organoid). Overall, we validated the capacity of FRC-QE to faithfully capture image quality in three-dimensions and show its ability to quantitatively measure image quality and thus clearing efficiency. We will next show how FRC-QE can be used across protocols and for different microscopy techniques, and finally show how FRC-QE is the only metric that enabled us to determine the best clearing protocol for our samples.

### Using FRC-QE across protocols

We compared FRC-QE across protocols using multi-view light-sheet microscopy data. Previously it has been shown that computationally fusing multiple images of the same object taken from multiple angles significantly increased the volume of the sample that can be imaged with high resolution (24, 25). Here, we acquired four volumes for each cleared organoid using dual-illumination (left and right sided light-sheet illumination) and acquisition from opposite acquisition angles (0 and 180 degrees), which were registered and fused using BigStitcher (16). FRC-QE faithfully captures the increase in image quality from individually fusing left and right illumination (Supplementary figure 3) as well as from additionally combining the opposite acquisition angles (Figure 3a-c). Consequently, organoids that were imaged with only a single angle show image quality that is constantly decreasing with imaging depth, resulting in one peak on the left or on the right side of the FRC-QE score plot. After multi-view fusion of all four volumes, FRC-QE showed as expected two clear peaks towards the edges of the imaged sample and a reduced quality in the center of the organoid. When comparing the ClearT2 and Fructose-Glycerol clearing protocols as performed on our samples, we noticed insufficient clearing at the center of the ClearT2-cleared organoid (Figure 3b, iii), while at the surface individual nuclei were clearly resolved (Figure 3b, iv). For the Fructose-Glycerol-cleared organoid, image quality was perceived slightly lower on the surface of the organoid but more stable throughout the organoid, indicating successful, homogeneous clearing. Both of these characteristics were captured by FRC-QE, resulting in a higher score at the surface for the ClearT2 organoid while the FRC-QE score for the Fructose-Glycerol sample did not decrease significantly towards the center (Figure 3c). Hence, the FRC-QE score does not only capture the expected improved resolution of multi-view datasets but also provides a metric that allows quantitative comparison between different clearing protocols and regions within the organoid.

**Fig. 3.**
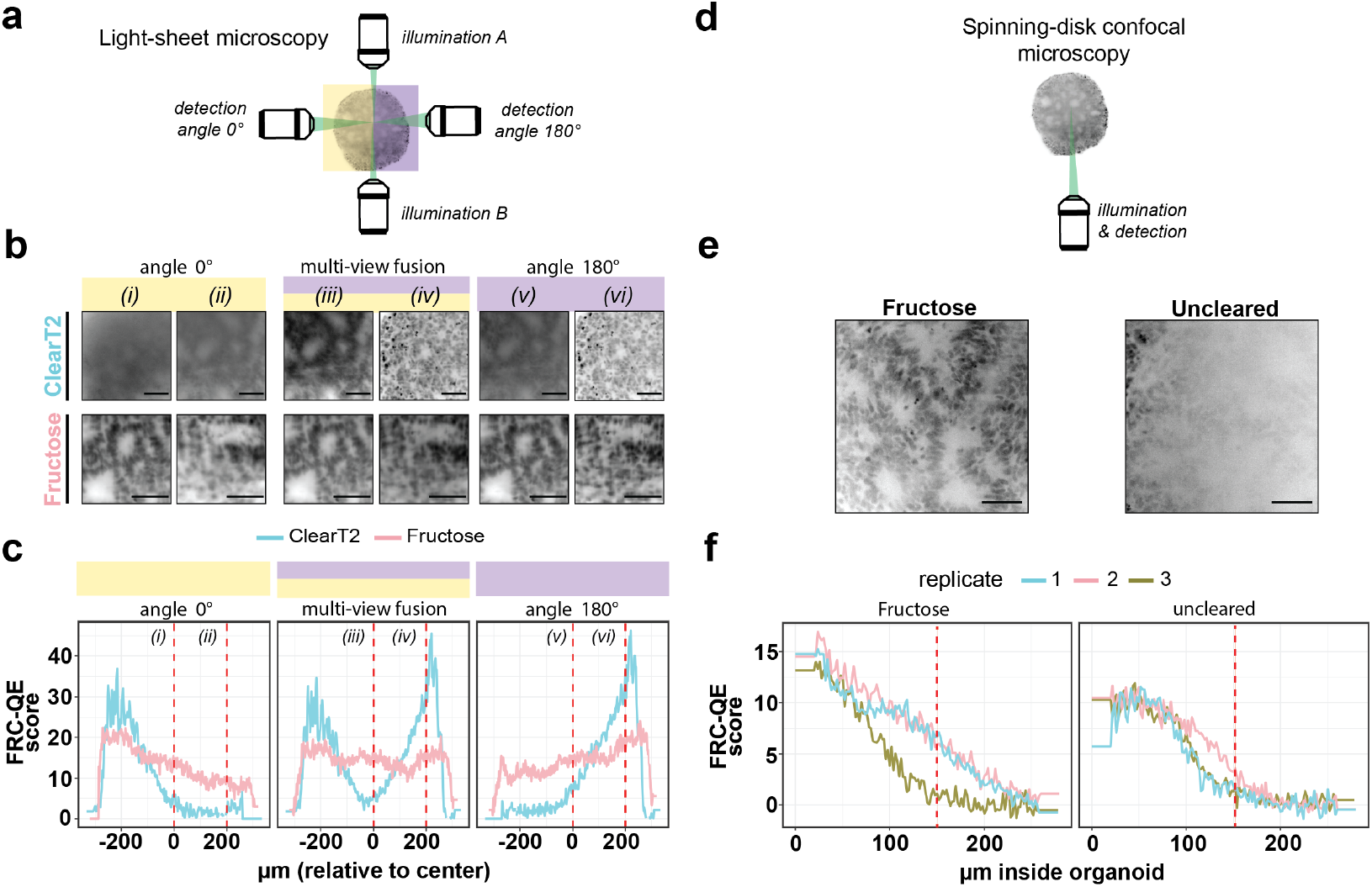
Using FRC-QE for different microscopy modalities. (a) Schematic of multi-view light-sheet microscopy. (b) Depicts example images of multi-view fused as well as left and right illumination only datasets for two different protocols. (c) Represents the corresponding FRC-QE score across the z-axis of the organoid. (d) Schematic of spinningdisk confocal microscopy. (e) Two example images for Fructose-Glycerol cleared and uncleared organoids (both 150 μm inside the organoid). (f) Depicts corresponding FRC-QE score curves for 3 replicates each. Scale bars correspond to 50 μm. Dotted red lines correspond to the corresponding panels above as indicated. Brightness and contrast was adjusted individually for the example images.

### Using FRC-QE for different microscopy modalities

To further validate the FRC-QE approach for different microscopy modalities, we compared Fructose-Glycerol-cleared organoids to uncleared organoids using a spinningdisk confocal microscope (Figure 3d). As expected, when comparing z-slices at the center of the organoid, images from uncleared controls did not offer sufficient resolution to distinguish individual nuclei, whereas in Fructose-Glycerol-cleared organoids individual nuclei could be identified (Figure 3e). This observation is recapitulated by FRC-QE, which results in a steeper slope in uncleared organoids. Importantly, the Fructose-Glycerol-cleared organoids contained one outlier showing an FRC-QE similar to uncleared organoids (Figure 3f, green). Manual inspection of this organoid confirmed unsuccessful clearing in that particular case (Supplementary figure 4, replicate 3). Thus, this observation underlines the capability of FRC-QE to robustly identify differences in clearing efficiency without the need for manual inspection of the image data. This will be particularly important when imaging a higher number of cleared samples (e.g. in the case of automated organoid screens (26)), where manual inspection of all samples for quality control is not feasible. Moreover, estimating clearing efficiency automatically over many replicates will also be important to determine the experimental robustness and reproducibility of a given method.

### FRC-QE based three-dimensional analysis helps to identify a suitable clearing protocol

After having validated that FRC-QE faithfully captured image quality across protocols and for different microscopy set-ups, we next sought to use this metric for identifying the clearing protocol that resulted in the best clearing result, given our samples, stainings, cost, and time effort. As expected, in all protocols nuclear structures could be easily visually identified at the surface of the organoid. However, image resolution differed at the center, with ClearT2 and ScaleA2 resulting in blurred objects compared to the Fructose-Glycerol protocol (Figure 4a-b), indicating differences in performance between protocols as we performed them. Measuring image contrast along the z-axis, it was possible to capture the observed protocol-dependent differences in clearing efficiency to some extent (Figure 4c). Nevertheless, the resulting curves remained unstable and made comparison between protocols difficult, especially when trying to interpret the highly variable sharp peaks at the surface of the organoid. Having validated the capability of DCT Shannon entropy to capture relative differences in resolution across a single organoid, we next tested its performance across protocols (Figure 4d). The metric resulted in equally low values for the center regions of all tested samples, only differing in entropy values for the respective surface regions. Thus, while faithfully capturing relative differences in clearing efficiency in one organoid (Figure 2g), we note that DCT Shannon entropy is not suited for comparison between samples since it did not recapitulate the visually apparent differences in clearing efficiency that we observed for the three protocols. In contrast, FRC-QE recapitulated clear discrepancies between the protocols as we observed them (Figure 4e). While the FRC-QE score of the other two tested protocols steadily decreased towards the center of the organoid, Fructose-Glycerol-cleared samples did preserve a constant FRC-QE score throughout the entire volumes of the cleared organoids. Hence, FRC-QE was the only algorithm that recapitulated the observed differences in image quality and could aid identify a clearing protocol which gave the best result in terms of signal-to-noise ratio and minimal light scattering. Consequently, quantifying DCT Shannon entropy across slices of all protocols and replicates did not give sufficient information to identify the most successfully cleared samples (Figure 4f). In comparison, the average FRC-QE score across z-slices was significantly higher in Fructose-Glycerol-cleared samples compared to ClearT2 or ScaleA2 clearing (Figure 4g), confirming the visually apparent differences in clearing efficiency (Figure 4a-b). In summary, FRC-QE provides a robust and comparable metric for clearing efficiency, providing objective guidance when trying to identify the most suitable clearing protocol for a given type of organoid sample.

**Fig. 4.**
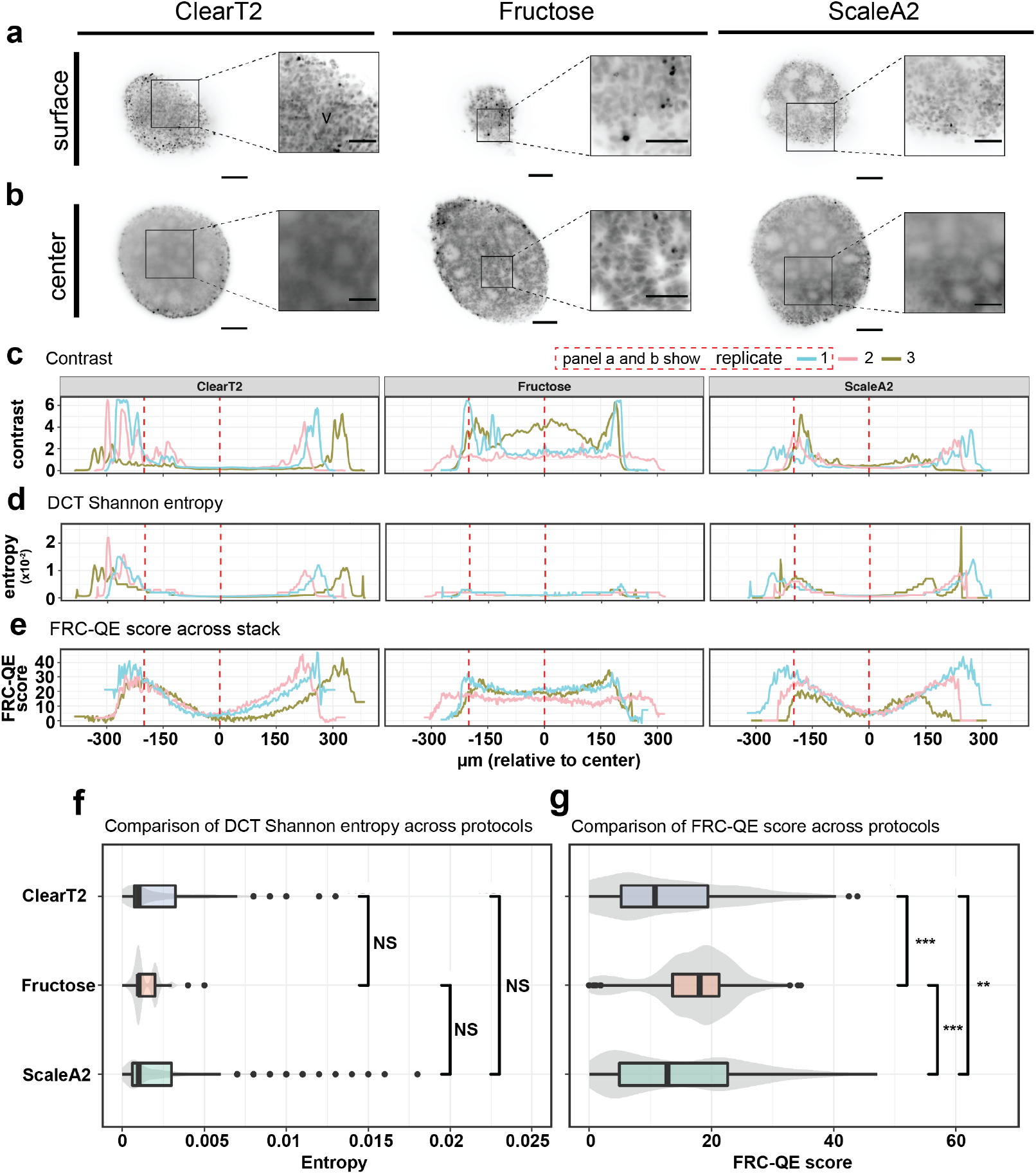
Comparing image quality across protocols using FRC-QE. (a-b) Example images for each protocol, where (a) is a location at the edge of the organoid (center location minus 200 μm) and (b) corresponds to the center of the organoid. (c-e) Image quality metrics across the organoid for three replicates each, all imaged with light-sheet microscopy and multi-view reconstructed. Replicate 1 (light blue) always corresponds to the example images shown above. Fructose-Glycerol clearing showed the lowest decrease in the FRC-QE score in the center of the organoid, indicating successful clearing. (f-g) Boxplots comparing image quality estimates across protocols. Each dataset was sampled to an equal number of measured slices (600 per protocol) and the same images were analysed by DCT Shannon entropy and FRC-QE, respectively for (f) and (g). Boxplot center line: median. Box limits: First and third quantiles. Grey shaded area: violin plot for the same dataset. Statistical significance values were calculated using Wilcoxon rank test, NS (not significant): p > 0.05, **: p < 0.01, ***: p < 0.001. Dotted red lines correspond to the corresponding image slices above as indicated. Scale bars correspond to 100 μm and 50 μm for large panels and inlets, respectively. Brightness and contrast was adjusted individually for the example images.

## Discussion

We introduce FRC-QE as a new metric to automatically assess clearing efficiency from three-dimensional fluorescence microscopy images. We first validated that FRC-QE reliably captured differences in image quality across single, whole organoids. FRC-QE performed with better accuracy than intensity or contrast measurements and similar to DCT Shannon entropy, a state-of-the-art autofocus algorithm (17, 18). Furthermore, FRC-QE can be applied to image data from different microscopy modalities and is comparable across protocols. We highlight this capability using FRC-QE calculation to identify the clearing protocol suiting best our samples. We believe that this is of particular importance for the clearing field. Until now, researchers trying to establish (or even develop) a new clearing protocol for a sample of interest were lacking an appropriate method that allowed assessing clearing efficiency automatically. However, such a method is crucial in order to optimize the protocol, adapt it to the challenges of the sample of interest and to assess variability between experimental replicates. To fill this gap, we provide the open-source FRC-QE ImgLib2-based (27) implementation as a stand-alone Fiji (28) plugin that is macro-scriptable and can therefore batch-process multiple images automatically. Therefore, it allows rapid testing for comparison of different clearing protocols and will also be helpful for finetuning specific parameters of a selected protocol to allow its optimization in a high throughput manner. While other quality metrics such as the DCT Shannon entropy perform well within a given sample or experiment, they were not developed for comparing between different experiments and therefore cannot be used as an absolute metric for assessing clearing efficiency across trials. It is thus important to emphasize that FRC-QE captures differences in clearing efficiency within one sample as well as across replicates and different protocols. Since the quality estimation is directly computed from the images, FRC-QE calculation can be performed on several fluorescent dyes or immunohistochemistry stainings of specific cells or subcellular structures that might differ in their compatibility with a given clearing protocol. Conceptually, FRC-QE is superior to methods like DCT Shannon entropy for comparing different samples because its measurements are more abstract. DCT Shannon entropy measures the information content of the frequency space. However, noise also produces high frequency components and DCT Shannon entropy is therefore not able to differentiate between noise and content. DCT Shannon entropy nevertheless works robustly on single stacks, which is required for autofocussing, since noise patterns typically do not change locally. FRC-QE on the other hand measures the correlation in between individual frequencies relative to a background correlation and is thereby able to differentiate noise from actual image content. Furthermore, it is able to ignore artifacts such as camera noise or local dirt. This allows for a higher degree of invariance and thereby enables the comparison of different clearing protocols for example. At the same time, it is important to note that the FRC-QE score represents arbitrary numbers that do not directly relate to an actual measurement of image resolution, but only allow for a relative comparison. Importantly, the FRC-QE score depends on the area in which it is computed, the z-spacing, type of image content (e.g. nuclear stain) and the point spread function (PSF). These parameters should therefore be held constant during a series of comparisons. When establishing a clearing protocol of choice for a new sample type in their laboratory, researchers are faced with a myriad of different protocols to choose from, each with specific advantages and drawbacks. Importantly, protocols will differ e.g. in terms of experimental duration, cost, ability to reproduce it in the lab, compatibility with the microscope of choice and the given sample. The most suitable protocol is therefore often the one that achieves the necessary image quality to gain a certain insight given the lowest effort and cost. Hence, by using the FRC-QE score which allows automated quality estimation across multiple samples, researchers will be able to reliably assess the obtained image quality and thereby clearing efficiency under different experimental conditions. Overall, we believe that image quality estimation using FRC-QE will facilitate and significantly ease the process of choosing the right clearing protocol for a given biological sample.

## Data availability

The source code is licensed under GPLv3 and is available on github together with documentation on how to install and use the Fiji plugin: https://github.com/PreibischLab/FRC-QE. All image datasets used in this study are available for download from http://bit.ly/FRC-QE_raw_data. All measured, raw data together with analysis code to reproduce Figure 2-4 are available under https://github.com/PreibischLab/FRC-QE/tree/master/analysis_scripts. Current versions of the plugin, a documentation as well as an example ImageJ macro for automated execution can be downloaded from the same GitHub repository.

## ACKNOWLEDGEMENTS

This work was supported by the Bundesministerium für Bildung und Forschung, Germany (BMBF, grant no. 031L0142A to S.P. and P.M., and in part by grant no. 16GW0191 to P.M. and H.S.) and HFSP grant RGP0021/2018-102 to S.P. and F.P. F.P. was funded by a PhD fellowship from Studienstiftung des deutschen Volkes. S.P. was funded by MDC Berlin. N.S. was funded by FAPESP’s program of Research Internships Abroad (BEPE, 2019/06305-4) during her visit to the Preibisch lab at the MDC Berlin. P.M. was supported by the BIH-Charité Clinical Scientist Program funded by Charité – Universitätsmedizin Berlin and the Berlin Institute of Health. E.T.C. was funded by the Hospital Sirio-Libanese/HSL/SBSHSL. Establishment of the Charité High Content Shared Facility was supported by the Deutsche Forschungsgemeinschaft (DFG, grant no. INST 335/591-1). We want to thank EU-Life and FAPESP (specifically Marta Dias Agostinho and Anamaria Camargo) for initiating this collaboration during the FAPESP/EU-LIFE Symposium on Cancer Genomics, Inflammation and Immunity 2016. We also thank Andrew Woehler for discussions and support during the project, Ricardo Henriques for sharing the bioRxiv LaTex template, and Loïc Royer for sharing the DCT Shannon entropy algorithm openly.

## AUTHOR CONTRIBUTIONS

N.S. and F.P. performed clearing and imaging; J.C. performed organoid culture, staining and confocal imaging; H.S. supervised organoid culture; S.P. and F.P. implemented the software; F.P. performed data analysis and drafted the figures; F.P., N.S., E.T.C, S.P. and P.M. conceived the project; S.P. and P.M. supervised the project; F.P., P.M. and S.P. wrote the manuscript. All authors read, edited, and approved the final version of the manuscript.

## Materials and Methods

### Cell Culture and generation of human cerebral organoids

Cerebral organoids were generated from the human induced pluripotent stem cell (hiPSC) line BIHi005-A (https://hpscreg.eu/cell-line/BIHi005-A). HiPSC line identity and integrity was verified at regular intervals. hiPSC were cultured in E8 medium in Geltrex-coated (Thermo Fisher) culture plates. For cerebral organoid induction, after single cell passaging hiPSC were first placed in neural induction medium (NIM, DMEM/F-12 (Thermo Fisher), 2.5 mM glutamine (Thermo Fisher), 15 mM HEPES (Thermo Fisher), 1x B27 (Thermo Fisher), 1x N2 (Thermo Fisher), 2 μM Dorsomorphin (Biovision), 10 μM SB431542 (Reagents Direct), 100 U/ml penicillin, 100 μg/ml streptomycin (Thermo Fisher)) for 6 days with daily medium changes. Next, medium was changed to neural expansion medium (NEM, 0.5x Neurobasal medium (Thermo Fisher), 0.5x Advanced DMEM/F12 (Thermo Fisher), 1x Neural Induction supplement (Thermo Fisher), 5 μM Y-27632 (Wako), 100 U/ml penicillin, 100 μg/ml streptomycin (Thermo Fisher)). Then, 7000 cells per well were seeded in a 96 well ultra low attachment round bottom plate (Corning) in neural expansion medium (NEM) and centrifuged (300x g, 5 minutes). Medium was changed to neural medium (NM, Neurobasal medium (Thermo Fisher), 1x B27 (Thermo Fisher), 2 mM Glutamax (Thermo Fisher), 20 ng/ml rhEGF (Peprotech), 20 ng/ml rhFGF-basic 154 a.a. (Peprotech), 100 U/ml penicillin, 100 μg/ml streptomycin (Thermo Fisher)), and replaced daily until day 4 and every other day thereafter. On day 6, medium was changed to neural differentiation medium (NDM, Neurobasal medium (Thermo Fisher), 1x B27 (Thermo Fisher), 2 mM Glutamax (Thermo Fisher), 20 ng/ml rhNT3 (Peprotech), 20 ng/ml rhBDNF (Peprotech), 100 U/ml penicillin, 100 μg/ml streptomycin (Thermo Fisher)). NDM was replaced every other day. Cerebral organoids were harvested on day 37.

### Sample clearing

For clearing, we used brain organoids of approximately 600 μm diameter. The organoids were stained with 5 μM Draq5, 5 μM Hoechst 33342 and 250 nM MitoTracker Red CMXRos (Thermo M7512) in NDM for 30 min at 37°C, fixed with 4% paraformaldehyde (PFA) for 30 min at room temperature, washed three times in PBS and stored in PBS. Clearing was performed based on three published clearing methods: ClearT2 (12), ScaleA2 (13) and Fructose-Glycerol (14) that were carried out according to the published protocols. Briefly, for the ClearT2 protocol, fixed organoids were incubated for 10 min at RT in a solution of 25% formamide/10% polyethylene glycol (PEG), followed by a 5 min incubation in a 50% formamide/20% PEG solution. Finally, organoids were immersed in fresh 50% formamide/20% PEG and incubated for 60 min at RT. All steps were carried out under gentle movement. In the ScaleA2 protocol, scale clearing solution consisted of 4 M urea, 0.1% wt/vol Triton X-100, and 10% wt/wt glycerol in water. Fixed organoids were incubated for 24h in fresh Scale clearing solution at room temperature and the solution was changed twice every 24 hours until 3 days. For the Fructose-glycerol solution, fixed organoids were placed on a heat block at 40°C, then resuspended in a 60% (vol/vol) glycerol and 2.5 M fructose/4% low melting point agarose solution. For light-sheet imaging, the mix was aspirated with a glass capillary and solidified at 4°C. Samples were incubated for 24h in the capillary before imaging. For spinning-disk confocal microscopy, fixed organoids were incubated for 20 minutes in fructose–glycerol clearing solution (60% (vol/vol) glycerol and 2.5M fructose) or PBS (negative control) before mounting.

### Light-sheet microscopy

For light-sheet imaging, cleared organoids were embedded in 2% low melting point agarose columns using glass capillaries (Zeiss). The light-sheet microscope used was a commercial Zeiss light-sheet Z.1 microscope. Glass capillaries were inserted into the imaging chamber of the microscope and the agarose column was extruded into the chamber filled with imaging solution. Chambers containing the organoids cleared by CleartT2 and ScaleA2 protocols were filled with water. For fructose-glycerol cleared organoids the imaging chamber was filled with fructose–glycerol clearing solution and we allowed the sample to settle in the imaging chamber overnight to improve sample clearing. Images were acquired using 10× illumina-tion objectives and a 20× detection objective. Draq5 was imaged with the 639 nm laser line. Laser power and microscope parameters are indicated in Supplementary table 1. Each organoid was imaged from two opposing angles, each with two illuminations, resulting in 4 views for each organoid. Multi-view reconstruction was performed in BigStitcher as previously described (16). Briefly, interest point-detection was performed on cell nuclei (Draq5 staining) for each view. Next, the 4 views were registered by the descriptor-based translation-invariant algorithm. Fused images were exported as TIFF files.

### Spinning-disk confocal microscopy

For spinning disk confocal microscopy, cleared organoids were placed in μ-Slide 8 Well chamberslides (Ibidi, 80827) and attached with one drop of 4% low melting point agarose. Imaging was performed on a PerkinElmer Opera Phenix with a 20x water objective (NA=1.0) in spinning-disk confocal mode, controlled by Harmony v 4 software. Lateral resolution for all images was 0.3 μm with 2 μm spacing between z-slices. Laser power and exposure time were kept constant for all samples.

### Image quality estimation algorithms

FRC-QE is based on the relative Fourier ring correlation (rFRC) that we previously developed (16). Briefly, we take advantage of the fact that consecutive image planes along the z-axis are very similar due to the axial extent of the PSF. Hence, computing the FRC between two z-slices and integrating it over all frequencies yields a robust quality metric, with low score indicating low image quality. However, we found that taking z-slices adjacent to each other can result in overall too high correlation in some areas of some image stacks. We hypothesize that inaccurate movement of the acquisition stage in z might be responsible. Instead, we therefore compute the FRC for slice z using the slices z+1 and z-1, which yields a more robust FRC readout (Supplementary Figure 2). To exclude artifacts caused by nonspecific patterned noise or imaging artifacts (e.g. induced by camera noise) leading to increased correlation at higher frequencies we calculate the relative FRC (rFRC) by subtracting a smoothed FRC baseline of z-slices spaced by z+m and z-m (default m=10) slices that are beyond the axial extent of the point spread function (PSF). The integral over the subtracted curves yields the FRC-QE score. FRC-QE does not represent an actual measure of image resolution. Instead it describes how much more image correlation by frequency there is at the location where it is computed, as compared slices that are out of range of the PSF. Naturally, the actual values of the FRC-QE score depend significantly on the z-step size, type of content (e.g. nuclei stain), the PSF size and the FFT size in which the FRC is computed. It is therefore important to keep these parameters constant when comparing outcomes of different experiments. We implemented the adapted rFRC calculation at defined block sizes for the FRC-QE Fiji plugin. For the light-sheet microscopy data shown here, we used a 400×400 pixel window (spanning all z-slices) as input for each organoid to compute the FRC-QE score using a 200×200 pixel block, resulting in 4 subtiles spanning the 400×400px window, each corresponding to a distinct FRC-QE score for each plane. For each plane, we take the median value of the 4 subtiles yielding the final FRC-QE score. Thus, by taking the median value of 4 spatially separated tiles, high frequency correlations caused by imaging artifacts are suppressed. We compare this value to average pixel intensity and contrast per plane. Contrast was calculated for each plane as minimum pixel intensity subtracted from maximum pixel intensity. All intensity and contrast values were divided by 10^4^ for better readability. Normalized discrete cosine transform (DCT) Shannon entropy was imported and called as previously described (17).

### Statistical analysis and visualization

Statistical analysis and visualization of image data was done in R (Version 3.6.1), using ggplot2 (29) and the dplyr (30) package.

## Supplementary figures

**Supplementary figure 1.**
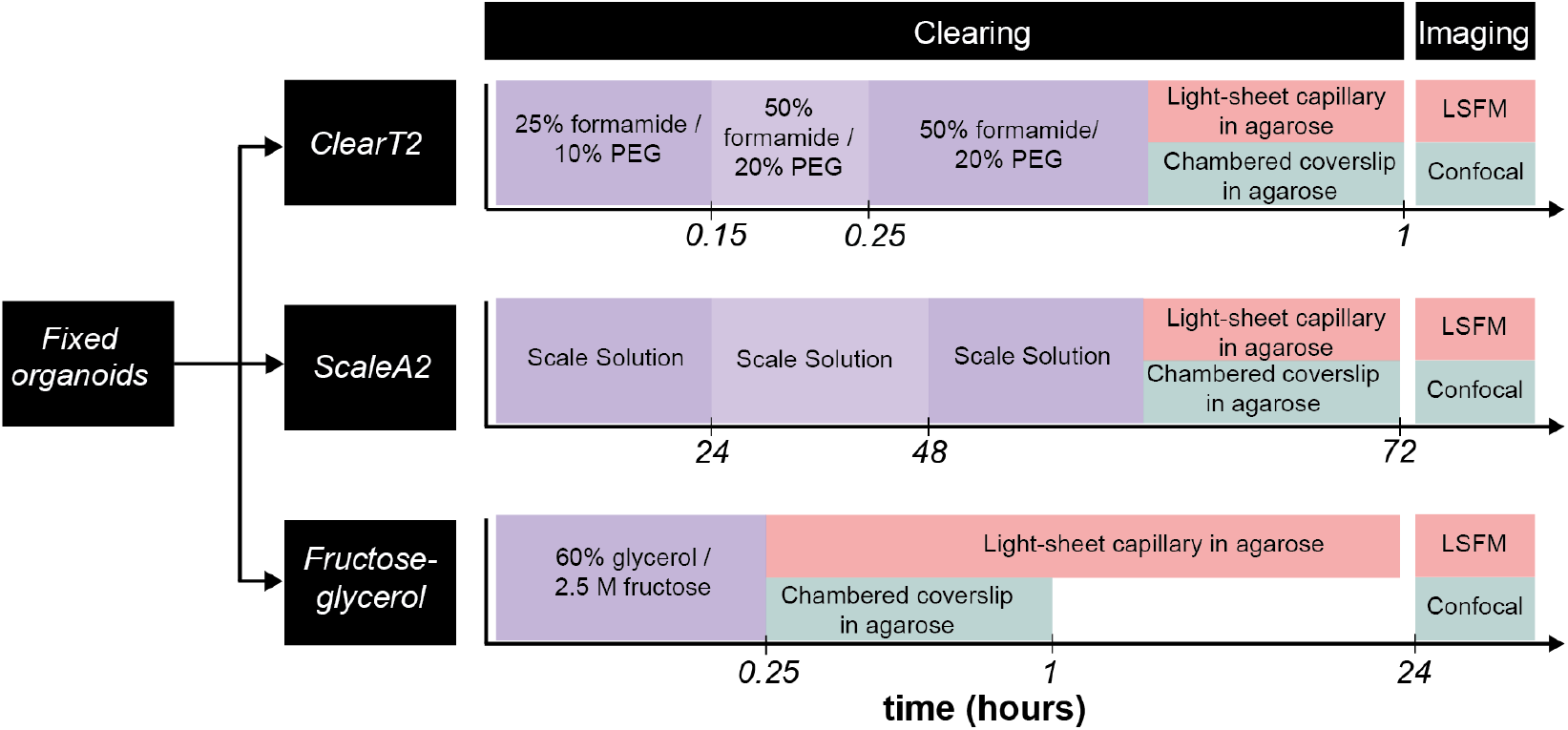
Overview scheme comparing tested clearing protocols and respective experimental timing (not to scale)

**Supplementary figure 2.**
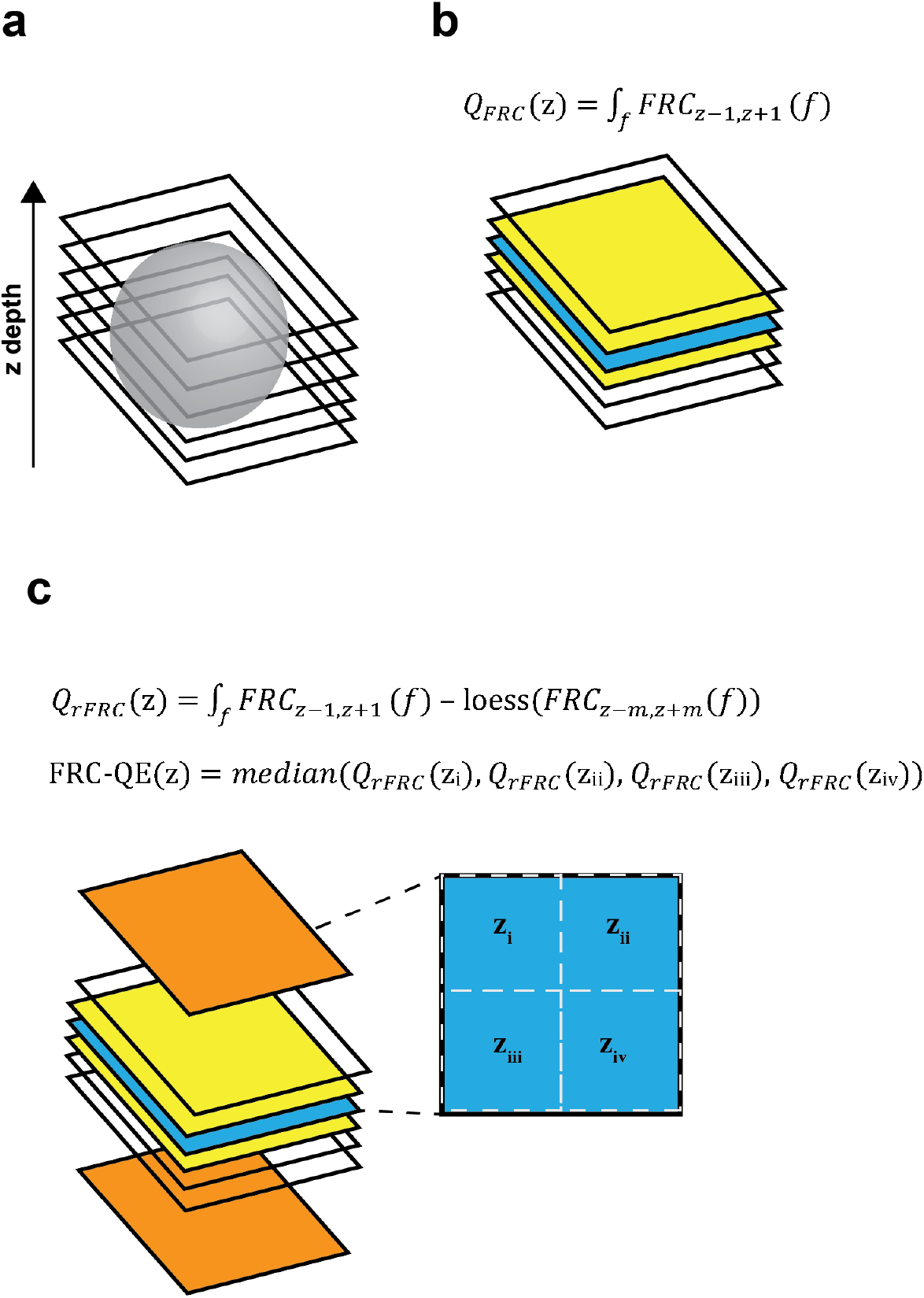
Overview of the FRC-QE implementation. (a) For three-dimensional imaging, we image an object (e.g. an organoid) by acquiring several adjacent image planes. (b) FRC describes the perspatial-frequency (f) correlation between two independent realizations. For each plane we correlate its adjacent planes (z-1 and z+1) taking advantage of the fact that they contain very similar information due to the axial extent of the PSF. (c) To exclude nonspecific patterned noise (e.g. camera noise), a smoothed baseline FRC of planes m slices away of z is subtracted from averaged correlation scores between adjacent planes. To further reduce the influence of imaging artifacts (e.g. bright dots) on the FRC-QE, the metric can be calculated blockwise (e.g. into 4 equally sized blocks spanning the field of view) and the final FRC score per slice is calculated as the median of all blocks.

**Supplementary figure 3.**
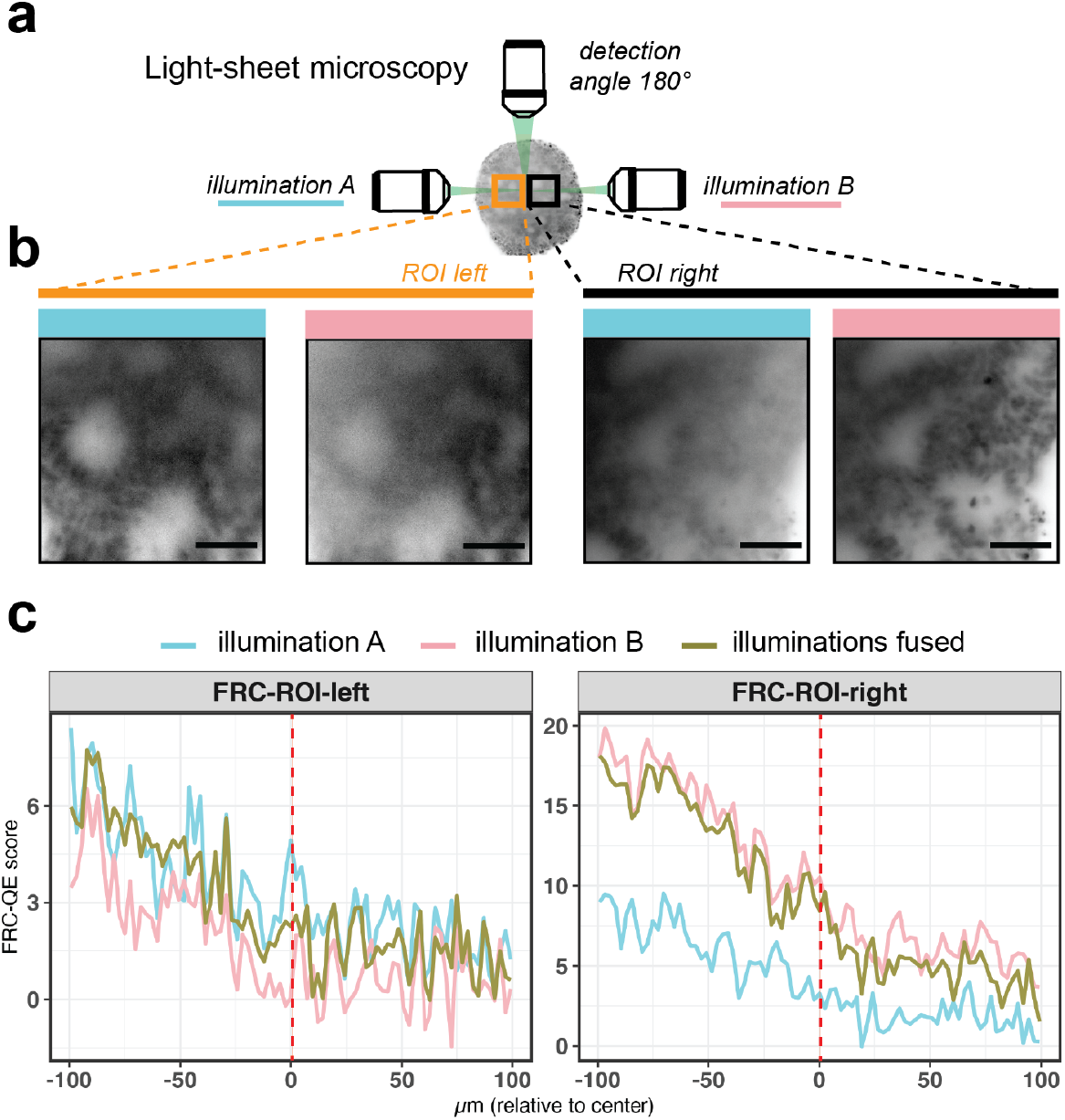
FRC-QE captures illumination-dependent differences in image quality. (a) Schematic of the two quantified regions of interest (ROI) (orange and black), being more close to one or the other illumination side. (b) example image (unfused, ClearT2 protocol, insufficient clearing) for each of the ROIs and illumination sides. (c) FRC-QE score for individual illumination sides as well as fused images for each ROI respectively. Dotted red lines correspond to the corresponding z-slice position of the image stacks shown in (b). Scale bars correspond to 50 μm.

**Supplementary figure 4.**
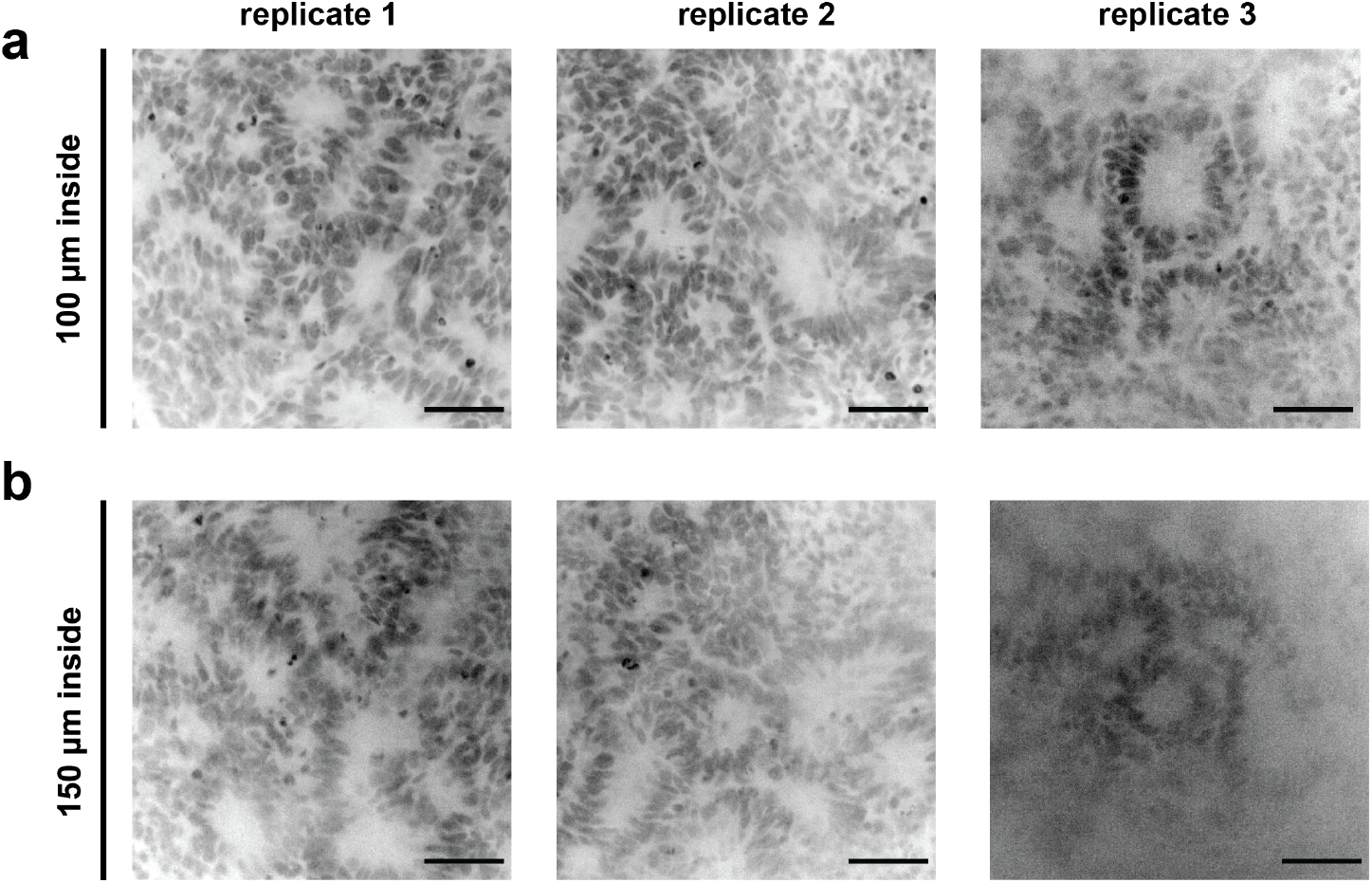
Differences in clearing efficiency for Fructose cleared organoids on a spinning-disk confocal microscope. (a) shows similar image quality for Draq5 signal in three replicates of Fructose cleared organoids at the surface of the organoid (50 μm). (b) Towards the center of the organoid (150 μm inside the organoid) image quality differs between replicates with replicate 3 showing lower image quality and increased light scattering.

**Supplementary table 1.** Imaging parameters for light-sheet imaging

| Protocol | ID | replicate ID in Figure 4 | x (μm) | y (μm) | z (μm) | lightsheet thickness (μm) | laser intensities | exposure time (ms) |
| --- | --- | --- | --- | --- | --- | --- | --- | --- |
| ClearT2 | org1 | replicate 3 | 0.458 | 0.458 | 2.42 | 4.8 | H: 20%, MT: 15%, D5: 15% | 30 |
|  | org2 | replicate 2 | 0.458 | 0.458 | 2.42 | 4.8 | H: 20%, MT: 15%, D5: 15% | 30 |
|  | org3 | replicate 1 | 0.458 | 0.458 | 2.42 | 4.8 | H: 20%, MT: 15%, D5: 15% | 30 |
| Fructose | org2 | replicate 3 | 0.318 | 0.318 | 1.7 | 5.61 | H: 20%, MT: 15%, D5: 15% | 20 |
|  | org4 | replicate 2 | 0.301 | 0.301 | 1.45 | 5.64 | H:10%, MT: 5%, D5: 10% | 30 |
|  | org5 | replicate 1 | 0.301 | 0.301 | 1.55 | 6.85 | H:10%, MT: 5%, D5: 10% | 30 |
| ScaleA2 | org1 | replicate 3 | 0.458 | 0.458 | 2.42 | 4.8 | H: 20%, MT: 15%, D5: 15% | 30 |
|  | org2 | replicate 1 | 0.458 | 0.458 | 2.42 | 4.8 | H: 20%, MT: 15%, D5: 15% | 30 |
|  | org3 | replicate 2 | 0.458 | 0.458 | 2.42 | 4.8 | H: 20%, MT: 15%, D5: 15% | 30 |

## Notes

https://github.com/PreibischLab/FRC-QE

### Competing Interest Statement

The authors have declared no competing interest.

## Bibliography

1. Jihoon Kim, Bon-Kyoung Koo, and Juergen A. Knoblich. Human organoids: model systems for human biology and medicine. Nature Reviews Molecular Cell Biology, 2020. doi: 10.1038/s41580-020-0259-3.

2. Frans Schutgens and Hans Clevers. Human Organoids: Tools for Understanding Biology and Treating Diseases. Annual Review of Pathology: Mechanisms of Disease, 15(1):211 – 234, 2020. doi: 10.1146/annurev-pathmechdis-012419-032611.

3. Werner Spalteholz. Über das Durchsichtigmachen von menschlichen und tierischen Präparaten und seine theoretischen Bedingungen, nebst Anhang: Über Knochenfärbung. Hirzel, 1914.

4. Douglas S. Richardson and Jeff W. Lichtman. Clarifying Tissue Clearing. Cell, 162(2): 246–257, 2015. doi: 10.1016/j.cell.2015.06.067.

5. Hiroki R. Ueda, Ali Ertürk, Kwanghun Chung, Viviana Gradinaru, Alain Chédotal, Pavel Tomancak, and Philipp J. Keller. Tissue clearing and its applications in neuroscience. Nature Reviews Neuroscience, 21(2):61–79, 2020. doi: 10.1038/s41583-019-0250-1.

6. Hiroki R. Ueda, Hans Ulrich Dodt, Pavel Osten, Michael N. Economo, Jayaram Chandrashekar, and Philipp J. Keller. Whole-Brain Profiling of Cells and Circuits in Mammals by Tissue Clearing and Light-Sheet Microscopy. Neuron, 106(3):369–387, 2020. doi: 10.1016/j.neuron.2020.03.004.

7. Paweł Matryba, Leszek Kaczmarek, and Jakub Golab. Advances in Ex Situ Tissue Optical Clearing, August 2019. ISSN: 1863-8899 Issue: 8 Publisher: John Wiley & Sons, Ltd Volume: 13.

8. Ruiyao Cai, Chenchen Pan, Alireza Ghasemigharagoz, Mihail Ivilinov Todorov, Benjamin Förstera, Shan Zhao, Harsharan S. Bhatia, Arnaldo Parra-Damas, Leander Mrowka, Delphine Theodorou, Markus Rempfler, Anna L. R. Xavier, Benjamin T. Kress, Corinne Benakis, Hanno Steinke, Sabine Liebscher, Ingo Bechmann, Arthur Liesz, Bjoern Menze, Martin Kerschensteiner, Maiken Nedergaard, and Ali Ertürk. Panoptic imaging of transparent mice reveals whole-body neuronal projections and skull–meninges connections. Nature Neuroscience, 22(2):317–327, 2019. doi: 10.1038/s41593-018-0301-3.

9. Peng Wan, Jingtan Zhu, Jianyi Xu, Yusha Li, Tingting Yu, and Dan Zhu. Evaluation of seven optical clearing methods in mouse brain. Neurophotonics, 5(03):1, 2018. doi: 10.1117/1.nph.5.3.035007.

10. Marko Pende, Karim Vadiwala, Hannah Schmidbaur, Alexander W. Stockinger, Prayag Murawala, Saiedeh Saghafi, Marcus P. S. Dekens, Klaus Becker, Roger Revilla-i Domingo, Sofia-Christina Papadopoulos, Martin Zurl, Pawel Pasierbek, Oleg Simakov, Elly M. Tanaka, Florian Raible, and Hans-Ulrich Dodt. A versatile depigmentation, clearing, and labeling method for exploring nervous system diversity. Science Advances, 6(22):eaba0365, 2020. doi: 10.1126/sciadv.aba0365.

11. Katherine N. Elfer, Andrew B. Sholl, Mei Wang, David B. Tulman, Sree H. Mandava, Benjamin R. Lee, and J. Quincy Brown. DRAQ5 and eosin (‘D&E’) as an analog to hematoxylin and eosin for rapid fluorescence histology of fresh tissues. PLoS ONE, 11(10):1–18, 2016. doi: 10.1371/journal.pone.0165530.

12. Takaaki Kuwajima, Austen A. Sitko, Punita Bhansali, Chris Jurgens, William Guido, and Carol Mason. ClearT: A detergent- and solvent-free clearing method for neuronal and nonneuronal tissue. Development (Cambridge), 140(6):1364–1368, 2013. doi: 10.1242/dev.091844.

13. Hiroshi Hama, Hiroshi Kurokawa, Hiroyuki Kawano, Ryoko Ando, Tomomi Shimogori, Hisayori Noda, Kiyoko Fukami, Asako Sakaue-Sawano, and Atsushi Miyawaki. Scale: A chemical approach for fluorescence imaging and reconstruction of transparent mouse brain. Nature Neuroscience, 14(11):1481–1488, 2011. doi: 10.1038/nn.2928.

14. Johanna F. Dekkers, Maria Alieva, Lianne M. Wellens, Hendrikus C. R. Ariese, Paul R. Jamieson, Annelotte M. Vonk, Gimano D. Amatngalim, Huili Hu, Koen C. Oost, Hugo J. G. Snippert, Jeffrey M. Beekman, Ellen J. Wehrens, Jane E. Visvader, Hans Clevers, and Anne C. Rios. High-resolution 3D imaging of fixed and cleared organoids. Nature Protocols, 2019. doi: 10.1038/s41596-019-0160-8.

15. Jan Huisken, Jim Swoger, Filippo Del Bene, Joachim Wittbrodt, and Ernst H K Stelzer. Optical sectioning deep inside live embryos by selective plane illumination microscopy. Science (New York, N.Y.), 305(5686):1007–9, August 2004. doi: 10.1126/science.1100035.

16. David Hörl, Fabio Rojas Rusak, Friedrich Preusser, Paul Tillberg, Nadine Randel, Raghav K Chhetri, Albert Cardona, Philipp J Keller, Hartmann Harz, Heinrich Leonhardt, Mathias Treier, and Stephan Preibisch. BigStitcher: reconstructing high-resolution image datasets of cleared and expanded samples. Nature Methods, 2019. doi: 10.1038/s41592-019-0501-0.

17. Loïc A Royer, William C Lemon, Raghav K Chhetri, Yinan Wan, Michael Coleman, Eugene W Myers, and Philipp J Keller. Adaptive light-sheet microscopy for long-term, high-resolution imaging in living organisms. Nature Biotechnology, 34(12):1267–1278, 2016. doi: 10.1038/nbt.3708.

18. Jiaye He and Jan Huisken. Image quality guided smart rotation improves coverage in microscopy. Nature Communications, 11(1):1–9, 2020. doi: 10.1038/s41467-019-13821-y.

19. W. O. Saxton and W. Baumeister. The correlation averaging of a regularly arranged bacterial cell envelope protein. Journal of Microscopy, 127(2):127–138, 1982. doi: 10.1111/j.1365-2818.1982.tb00405.x.

20. Marin Van Heel. Similarity measures between images. Ultramicroscopy, 21(1):95 – 100, 1987. doi: https://doi.org/10.1016/0304-3991(87)90010-6.

21. Robert P.J. Nieuwenhuizen, Keith A. Lidke, Mark Bates, Daniela Leyton Puig, David Grünwald, Sjoerd Stallinga, and Bernd Rieger. Measuring image resolution in optical nanoscopy. Nature Methods, 10(6):557–562, 2013. doi: 10.1038/nmeth.2448.

22. Sami Koho, Giorgio Tortarolo, Marco Castello, Takahiro Deguchi, Alberto Diaspro, and Giuseppe Vicidomini. Fourier ring correlation simplifies image restoration in fluorescence microscopy. Nature Communications, 10(1), 2019. doi: 10.1038/s41467-019-11024-z.

23. A. Descloux, K. S. Grußmayer, and A. Radenovic. Parameter-free image resolution estimation based on decorrelation analysis. Nature Methods, 16(9):918–924, 2019. doi: 10.1038/s41592-019-0515-7.

24. Jim Swoger, Peter Verveer, Klaus Greger, Jan Huisken, and Ernst H.K. Stelzer. Multi-view image fusion improves resolution in three-dimensional microscopy. Optics Express, 15(13): 8029, 2007. doi: 10.1364/OE.15.008029.

25. Stephan Preibisch, Stephan Saalfeld, Johannes Schindelin, and Pavel Tomancak. Software for bead-based registration of selective plane illumination microscopy data. Nature methods, 7(6):418–9, 2010. doi: 10.1038/nmeth0610-418.

26. Nathalie Brandenberg, Sylke Hoehnel, Fabien Kuttler, Krisztian Homicsko, Camilla Ceroni, Till Ringel, Nikolce Gjorevski, Gerald Schwank, George Coukos, Gerardo Turcatti, and Matthias P. Lutolf. High-throughput automated organoid culture via stem-cell aggregation in microcavity arrays. Nature Biomedical Engineering, 2020. doi: 10.1038/s41551-020-0565-2.

27. Tobias Pietzsch, Stephan Preibisch, Pavel Tomančák, and Stephan Saalfeld. ImgLib2—generic image processing in Java. Bioinformatics, 28(22):3009–3011, November 2012. doi: 10.1093/bioinformatics/bts543.

28. Johannes Schindelin, Ignacio Arganda-Carreras, Erwin Frise, Verena Kaynig, Mark Longair, Tobias Pietzsch, Stephan Preibisch, Curtis Rueden, Stephan Saalfeld, Benjamin Schmid, Jean Yves Tinevez, Daniel James White, Volker Hartenstein, Kevin Eliceiri, Pavel Tomancak, and Albert Cardona. Fiji: An open-source platform for biological-image analysis. Nature Methods, 9(7):676–682, 2012. doi: 10.1038/nmeth.2019.

29. Hadley Wickham. ggplot2: Elegant Graphics for Data Analysis. 2016.

30. Hadley Wickham, Romain François, Lionel Henry, and Kirill Müller, dplyr: A Grammar of Data Manipulation. 2018.

